# Transcriptional stress induces the generation of DoGs in cancer cells

**DOI:** 10.1101/2023.11.15.567248

**Authors:** Francisco Ríos, Maritere Urióstegui-Arcos, Mario Zurita

## Abstract

A characteristic of the cellular response to stress, is the production of RNAs generated from readthrough transcription of genes, called downstream-of-<u>g</u>ene-(DoG) containing transcripts. Additionally, transcription inhibitor drugs are candidates for fighting cancer. In this work, we report the results of a bioinformatic analysis showing that one of the responses to transcription inhibition is the generation of DoGs in cancer cells. Although some genes that form DoGs were shared between the two cancer lines, there did not appear to be a functional correlation between them. However, our findings show that DoGs are generated as part of the cellular response to transcription inhibition like other types of cellular stress, suggesting that it may be part of the defense against transcriptional stress.

---

In recent years, different drugs, both traditional and new-generation drugs, that affect components of the basal transcription machinery, particularly those mediated by RNA polymerase II (RNA pol II), have been proposed for use against cancer [Filippakopoulos et al., 2010; Bradner et al., 2017; Villicaña, et al., 2014]. In fact, some of these substances have been tested in clinical trials [Senderowicz et al., 1998; Greeno et al., 2015; Martin et al., 2020; Noel et al., 2019].

The targets of these substances are mainly components of the preinitiation complex (PIC) or factors that are associated with the PIC either during transcription initiation or during transcription elongation [Kwiatkowski et al., 2014; Vervoort et al., 2022]. Among these drugs are substances that preferentially kill cancer cells, for example, inhibitors of RNA pol II itself, such as actinomycin D and α-amanitin [Cruz-Ruiz et al., 2021]; of the transcription and DNA repair complex TFIIH, such as triptolide (TPL) and THZ1; of the PTEFb complex such as flavopiridol; and others that affect transcription [Titov et al., 2011; Villicaña et al., 2013; Shu et al., 2016; Cao and Shilatifard 2014]. Additionally, compounds have been developed that affect the functions of the elongation machinery, for example, JQ1, which inhibits BRD4 and has a strong antiproliferative effect on different cancer cells [Filippakopoulos et al., 2010; Schwalm and Knapp, 2022].

Recently, we reported that when breast cancer cells are treated with TPL and THZ1, although the transcription of a large number of genes is inhibited, some genes are overexpressed [Uriostegui-Arcos et al., 2020]. In addition, the inhibition of TFIIH, as well as the inhibition of transcription initiation and elongation in other types of cancer cells using other substances, also induces the overexpression of a significant number of genes, some of which are shared among the responses to these substances [Cruz-Ruiz et al., 2021]. It is obvious that since cancer cells are exposed to a stress condition that inhibits transcription, they respond by overexpressing some genes, and we have named this as the transcriptional stress response (TSR) [Uriostegui-Arcos et all., 2020]. Interestingly, the reduction in the expression of some of these transcriptional stress response genes enhances the effect of TPL [Uriostegui-Arcos et all., 2020]. Therefore, TSR is another type of cellular stress response that includes overexpression of specific genes.

A characteristic that has recently been noted in cellular responses to different types of stress, such as osmotic stress and heat shock, as well as in some tumor cells and during viral infections, is the production of long noncoding RNAs (lncRNAs) generated from readthrough transcription of genes transcribed by RNA pol II; in other words, these transcripts are not processed at the site where these events normally occur downstream of the polyadenylation site (PAS) [Cardiello et al., 2018; Cugusi et al., 2022; Heinz et al., 2018; Morgan et al., 2022; Nemeroff et al., 1998; Rosa-Mercado et al., 2021; Shalgi et al., 2014; Vilborg et al., 2015]. These lncRNAs have been called downstream-of-gene (DoG)-containing transcripts. In general, the transcription of these DoGs is initiated in the promoter of a gene transcribed by RNA pol II. A DoG transcript has a minimum length of 5 kb and begins at the transcription termination site in the 3’ of the gene, although the length of this type of RNA can be different among genes, cellular stresses and even cell types. DoGs are retained in the nucleus, and it is likely that they remain associated with chromatin [Hennig et al., 2018; Morgan et al., 2022; Rosa-Mercado and Steitz 2022]. It has been documented that alterations in the transcription termination machinery, such as the cleavage and polyadenylation (CPA) complex or the XRN2 exonuclease, can result in the generation of DoGs [Rosa-Mercado and Steitz 2022; Eaton et al., 2020]. Likewise, it has been reported that hyperosmotic stress causes disruption of the Int11 subunit of the Integrator complex, which is an endonuclease that participates in terminating the transcription of some genes, triggering the production of DoGs [Rosa-Mercado et al., 2021]. Currently, it is unknown whether these lncRNAs have a role in the stress response or are simply the product of transcriptional dysregulation, which intriguingly preferentially affects some but not all genes in the genome.

With this information as a basis, we formulated the question of whether transcriptional stress, similar to other types of stress, also results in the generation of DoGs. To answer this question, we analyzed our transcriptome data from breast cancer cells treated with TPL [Uriostegui-Arcos et al., 2020], which inhibits transcription initiation, as well as public transcriptome data from other types of cancer cells treated with TPL and THZ1, which also inhibit transcription. In general, we found that TSR also induces the generation of DoGs, and an important proportion of these DoGs are shared between cell types. This confirms that inhibition of transcription induces a response similar to that induced by other types of stress and that it may be part of a mechanism to protect cancer cells against the effects of substances that inhibit transcription.

## Results

### Inhibition of transcription by TPL induces the formation of DoGs in breast cancer cells

TPL affects the initiation of transcription mediated by RNA pol II, inhibiting the ATPase activity of the XPB subunit of TFIIH, which is essential for the formation of the open complex, and induces the dissociation of the XPB-p52-p8 module from TFIIH [Uriostegui-Arcos et al., 2020]. As mentioned above, although TPL inhibits transcription, some genes are overexpressed [Uriostegui-Arcos et al., 2020]. While analyzing the transcripts by visualization of RNA-seq data obtained during the TSR in the MCF10-Er-Src transformed cell line, we found that some genes contained reads extending for several kb beyond the 3’ end. Based on this observation, we decided to analyze whether the TSR, similar to other types of cellular stress, induces the formation of DoGs. To this end, we analyzed the RNA-seq data of MCF10A-Er-Src cells with and without TPL treatment using ARTDeco [Roth et al., 2020], which have been used by several groups to identify DoGs from RNA-seq data. The minimum length of DoGs is generally considered to be approximately 5 kb; however, this is an arbitrary consideration. We decided to consider a minimum length of 4 kb of DoGs, since when comparing many genes in the presence or absence of transcriptional stress, it is obvious that many genes have a clear readthrough extension of approximately 4 kb when transcription is inhibited. For example, in the case of the breast cancer cells, we detected 374 DoGs with extension between 4000 and 5000 nucleotides (as show in Sup. Fig. 1).

The RNA-seq data from the breast cancer cells was single-end and non-stranded, and it may generate false positives using ARTDeco, as the authors mention in the program’s documentation [Roth et al., 2020]. Therefore, to use the RNA-seq data from non-stranded breast cancer cells and ARTDeco, we take advantage of RNA pol II ChIP-seq data from the same cells under the same conditions. It is known that in most genes that are transcriptionally active, RNA pol II pauses after synthesizing between 20 and 120 nt. Taking this data into account, we filtered the genes that produce DoGs by analyzing the presence of the RNA pol II peak in the 5’ region of the gene. In addition, another filter was applied by excluding those DoGs with RNA pol II peaks that maps in their DoG region (see material and methods). Thus, we were able to use the DoGs obtained by ARTDeco with confidence only from those genes that are transcriptionally active.

An example of a gene that generates DoGs in response to transcription inhibition in MCF10A-Er-Src cells is shown in Figure 1A. ARTDeco identified 789 DoGs in TPL-treated cells (Fig. 1B; Sup. Table 1). Although 326 DoGs were identified in untreated cells, the number was lower than in the cells treated with TPL (Fig. 1B). We observed that 183 of the DoGs identified in TPL-treated cells were shared with cells only incubated with DMSO (Fig. 1C). To know the expression levels of the DoGs for each condition, we performed a statistical analysis using ARTDeco ‘diff_exp_dogs’ mode (that use DESeq2), obtaining the log2 fold change and P-values from DoGs comparing the expression in TPL treated cells to the expression in untreated cells. Of the 932 DoGs identified in cells treated with TPL and with DMSO, 651 have a P-value ≤ 0.05. This analysis is represented in a heat map (Figure 1D), that shows two areas, one from untreated cells (blue area) and another one from TPL treated cells (red area), which indicates that the levels of downstream transcripts identified by ARTDeco in the cells treated with TPL are significantly higher than in the control cells (Fig 1D; Sup. Table 1. Similar criteria were used for the analysis of pancreatic cells (see ahead).

**Fig. 1.**
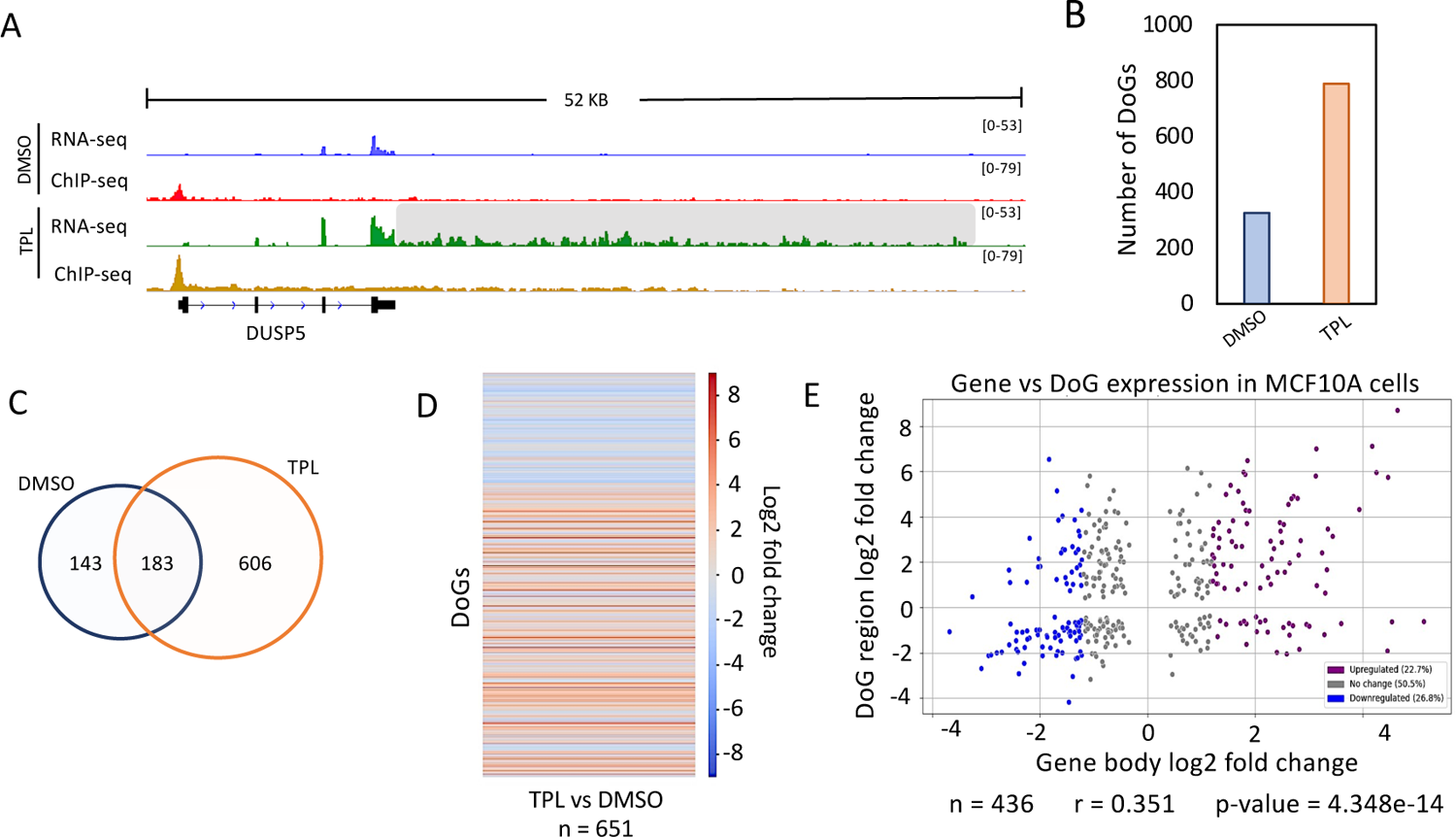
Inhibition of the XPB subunit of TFIIH by TPL induces the formation of DoGs in breast cancer cells. A) Browser image of RNA-seq and ChIP-seq profile of RNA pol II from MCF10A-Er-Src cells treated with DMSO or TPL. An example of a DoG-producing gene (*DUSP5*) identified with ARTDeco is presented, and the DoG region is delineated by a gray shadow (log_2_FC = 5.11 and p-value = 9.65e-71). B) Quantification of DoGs in MCF10A-Er-Src cells with DMSO and TPL. C) Venn diagram showing that 183 DoG-producing genes are shared between breast cancer cells treated with TPL and DMSO. D) Heatmap that exhibits the log_2_FC for each DoG from 651 DoGs found with TPL and DMSO with p-value ≤ 0.05. E) Scatter plot displaying the log_2_FC for the transcription levels of the DoG-producing genes (p-value ≤ 0.05) on the x-axis and the log_2_FC for the corresponding DoGs (p-value ≤ 0.05) on the y-axis. DoG-producing genes were classified as “Upregulated” (22.7%; purple dots) with log_2_FC ≥ 1.2, “Downregulated” (26.8%; blue dots) with log_2_FC ≤ −1.2, and “No change” (50.5%; gray dots) with log_2_FC between 1.2 and −1.2. Percentages for each category are displayed in the legend box. Pearson’s correlation coefficient and p-value were calculated, and both are shown below the graph.

In response to transcriptional stress, some genes are overexpressed. To determine if there is a correlation between genes that generate dogs and genes over-transcribed in response to TPL, we performed a scatter plot analysis of the expression of each DoG vs the expression of the corresponding gene (using the ARTDeco ‘diff_exp_read_in’ mode to obtain the log2 fold change of DoG-producing genes) and calculated the Pearson’s correlation coefficient with the gene’s transcripts that have a P-value ≤ 0.05 both in the DoG region and in the body of the gene (n=436) (Fig. 1E, Sup Table1).

Interestingly, there were no correlation between the expression levels of genes that are overexpressed in response to transcriptional stress and those from which DoGs are generated (Fig. 1E) [Uriostegui-Arcos et al., 2020]. This indicates that the generation of DoGs is not due to the overexpression of a gene, at least not in all cases. Taken together, the data analyzed in this section demonstrate that DoGs are generated as part of the response to transcriptional stress in breast cancer cells.

### Transcriptional stress also induces the generation of DoGs in pancreatic cancer cells

To investigate whether DoGs are generated in other types of cancer cells treated with TPL to inhibit transcription, we analyzed TPL-treated cells derived from pancreatic cancer in public RNA-seq data [Noel, et al., 2020]. Similar to our observations in MCF10A-Er-Src cells, we detected the presence of DoGs in these pancreatic cells following TPL treatment. An example of a gene generating a DoG is shown in Fig. 2A, which corresponds to the same gene depicted in Figure 1A, demonstrating a similar DoG formation in pancreatic cells. Intriguingly, as in MCF10A-Er-Src cells, untreated pancreatic cancer cells also displayed DoGs formation (Fig. 2B). However, treatment with TPL induced the generation of DoGs from a larger number of genes in this cell line (Fig. 2B; Sup. Table 2). When comparing the DoGs in DMSO-treated and TPL-treated pancreatic cancer cells, we found that 1406 DoGs were shared. And when those DoGs were excluded from the TPL treated cells, 2152 were found to be generated in response to transcriptional stress. This number was higher than that of DoGs in the MCF10A-Er-Src transcriptome and was due to differences in the sequencing depth in each experiment. Although the number of DoGs detected by ARTDeco was different between the two cell lines, out of the 606 found in breast cancer cells, 224 were also present in pancreatic cells (Fig. 2C; Sup. Table 2).

**Fig. 2.**
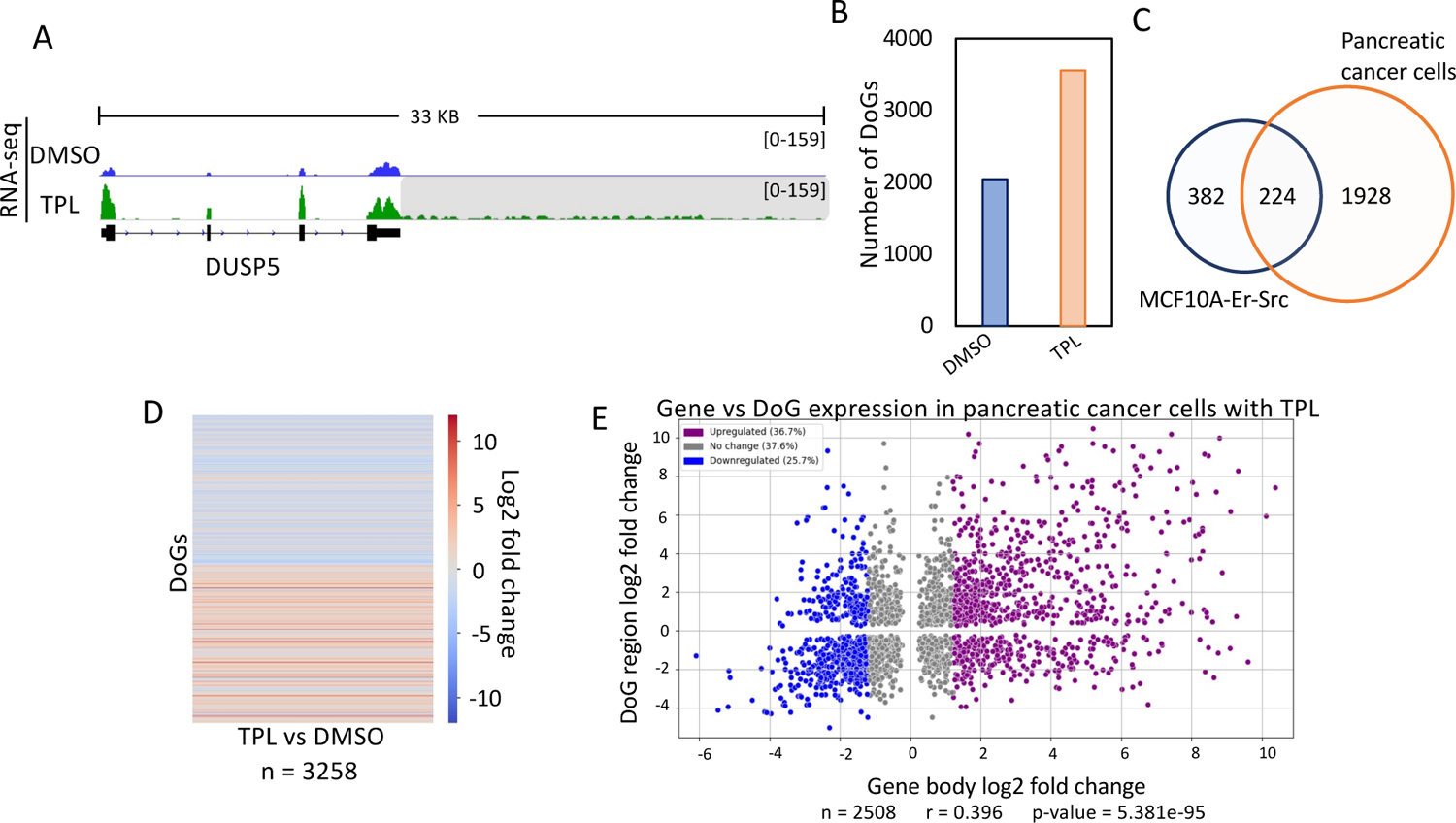
Inhibition of transcription by TPL in pancreatic cancer cells induces the formation of DoGs. A) Browser image of RNA-seq from pancreatic cancer cells treated with DMSO or TPL. The same example of a DoG-producing gene (*DUSP5*) also found in breast cancer cells in Fig. 1A is shown, and the DoG region is indicated by gray shadow (log_2_FC = 3.71 and p-value = 1.52e-27). B) Number of genes in pancreatic cancer cells in which there is the formation of DoGs in cells treated with DMSO and in response to the inhibition of transcription by TPL. C) Venn diagram showing that of the 606 genes that generate DoGs in breast cancer cells specific for the response to TPL, about 37% (224 DoGs) were also found in pancreatic cancer cells in response to TPL. D) Heatmap that exhibits the log_2_FC for each DoG from the 3258 DoGs found with TPL and DMSO in pancreatic cancer cells with p-value ≤ 0.05. E) Scatter plot displaying the log_2_FC for the transcription levels of the DoG-producing genes (p-value ≤ 0.05) on the x-axis and the log_2_FC for the corresponding DoGs (p-value ≤ 0.05) on the y-axis. DoG-producing genes were classified as “Upregulated” (36.7%; purple dots) with log_2_FC ≥ 1.2, “Downregulated” (25.7%; blue dots) with log_2_FC ≤ −1.2, and “No change” (37.6%; gray dots) with log_2_FC between 1.2 and −1.2. Percentages for each category are displayed in the legend box. Pearson’s correlation coefficient and p-value were calculated, and both are shown below the graph.

Similarly, the number of reads for the transcripts identified as DoGs generated by the TPL treatment in the pancreatic cells was quantified, and the corresponding values were determined and compared in a heatmap with the cells treated with DMSO, showing a clear increase in RNAs downstream of the identified genes (Fig. 2D, Sup. Table 2).

Additionally, as with breast cancer cells, there did not appear to be a correlation between genes with transcriptional upregulation in response to transcription inhibition and those that generate DoGs (Fig. 2E). In summary, the analysis presented in this section shows that transcription inhibition results in the generation of DoGs not only in the MCF10A-Er-Src cell line but also in other cancer cell types.

### Treatment of pancreatic cancer cells with THZ1 also induces the formation of DoGs

THZ1 is an inhibitor of the kinase activity of the Cdk7 subunit of TFIIH. Therefore, it is also an inhibitor of transcription mediated by RNA pol II and is a substance that is under investigation for the treatment of cancer [Kwiatkowski et al., 2014; Li et al 2019]. Therefore, we decided to explore whether Cdk7 inhibition in cancer cells also induces the formation of DoGs. Again, we used public RNA-seq data from a panel of pancreatic cancer cells [Noel et al., 2020] and ARTDeco. As expected, we found an increase of DoGs in the cells treated with THZ1. An example of a gene with its DoG is shown in figure 3A. As mentioned before, DoGs were also identified in untreated cells. However, we observed that the number of DoGs found by ARTDeco in THZ1 treated cells (3018 DoGs) was much larger than those in untreated cells (Fig. 3B; Sup Table 3). Again, as we did above, the DoGs found in THZ1 were compared to those from untreated cells, observing that 1049 DoGs were shared; leaving 1969 DoGs that are exclusively expressed in pancreatic cancer cells treated with THZ1.

**Fig. 3.**
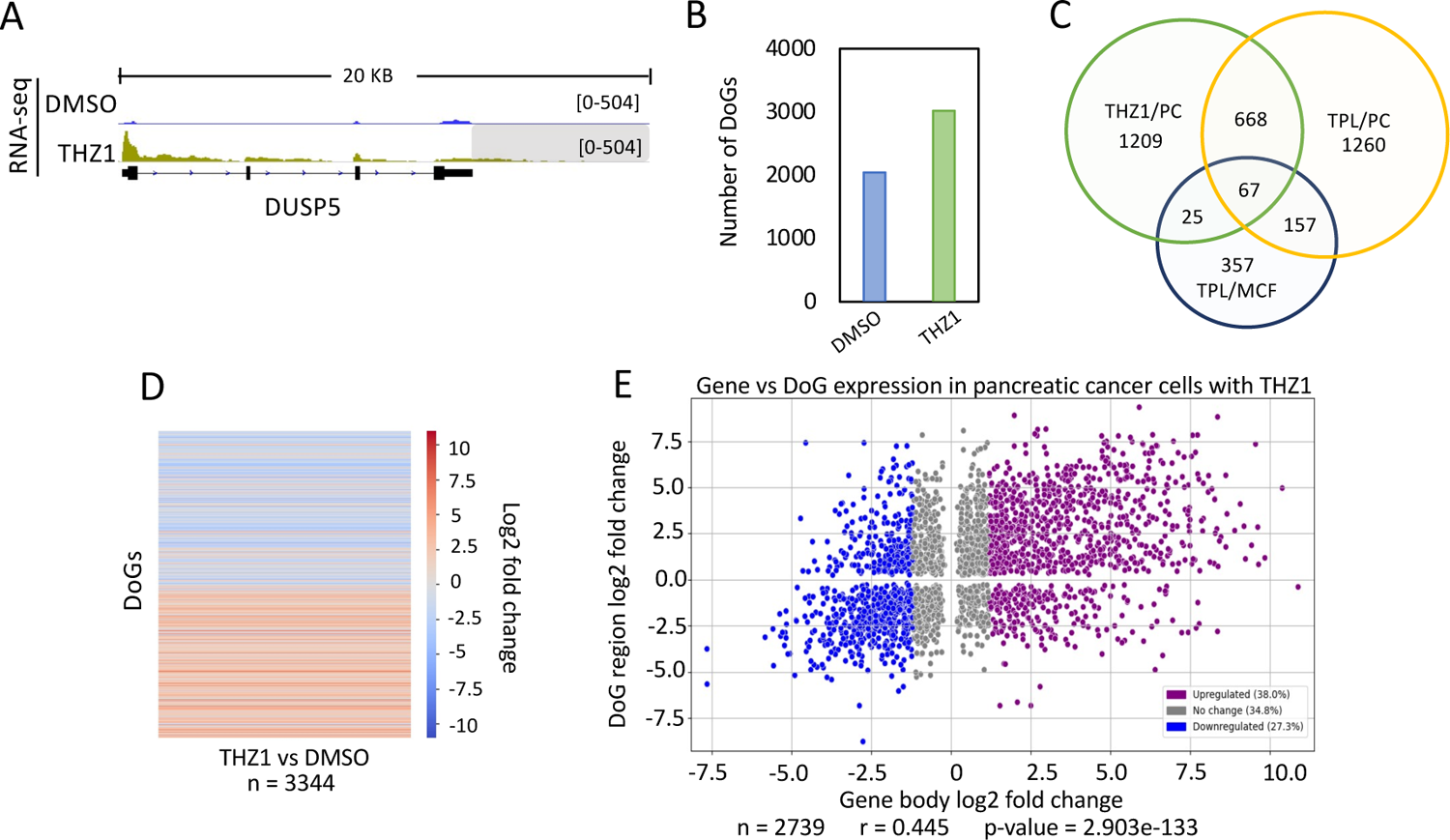
The inhibition of transcription with THZ1 induces the production of DoGs in pancreatic cancer cells. A) Browser image of RNA-seq from pancreatic cancer cells treated with DMSO or THZ1 showing the *DUSP5* gene that also undergoes the generation of a DoG in response to THZ1. DoG region is indicated by gray shadow (log_2_FC = 3.83 and p-value = 3.06e-29). B) Number of genes that generate DoGs in response to DMSO and THZ1 in pancreatic cancer cells. C) Venn diagram showing the genes that are shared between pancreatic cancer cells treated with TPL and THZ1 and with breast cancer cells treated with TPL. D) Heatmap showing the log_2_FC of downstream readthrough transcripts (p-value ≤ 0.05) between pancreatic cancer cells treated with THZ1 and DMSO. E) Scatterplot displaying the log_2_FC for the transcript levels of the DoG-producing genes (p-value ≤ 0.05) on the x-axis and the log_2_FC for the corresponding DoGs (with p-value ≤ 0.05) on the y-axis. DoG-producing genes were classified as “Upregulated” (38.0%; purple dots) with log_2_FC ≥ 1.2, “Downregulated” (27.3%; blue dots) with log_2_FC ≤ −1.2, and “No change” (34.8%; gray dots) with log_2_FC between 1.2 and −1.2. Percentages for each category are displayed in the legend box. Pearson’s correlation coefficient and p-value were calculated, and both are shown below the graph.

From this analysis, we found that a large number of DoGs generated in response to TPL and THZ1 treatment were shared between breast cancer and pancreatic cancer cells (Fig. 3C; Sup. Table 4). Intriguingly, even that both substances affect two different activities of TFIIH, there are DoGs that were unique for each treatment, even that the experiments were performed in the same cell type. However, among the genes detected in breast cancer cells in response to TPL, 92 were also detected in pancreatic cancer cells treated with THZ1, and 67 were shared between the pancreatic cells treated with THZ1 and TPL, as well as the breast cancer cells treated with TPL (Fig. 3C). This suggests that certain genes, when the activity of TFIIH is inhibited, are more likely to exhibit improper transcription termination in different cancer cells.

Once again, the difference in expression levels of DoGs was obtained and a heatmap showing the log2 fold change was plotted (Fig. 3D). The heat map shows that DoGs exclusively found in THZ1 treated cells have a significant higher expression in those cells than in untreated cells.

Taking in to account the expression levels of the DoG-producing genes and DoGs from THZ1 and DMSO treated cells (with p-value ≤ 0.05 in both the gene and the DoG region), we represented it in a scatter plot (Fig 3E); showing again that the expression level from the DoG-producing gene doesn’t correlate with the expression level of the DoG.

### Most of the DoGs generated in response to transcriptional stress do not overlap with the DoGs generated in response to by osmotic stress

The observation that some DoGs are generated in both breast and pancreatic cancer cells in response to transcriptional stress raises the question of whether these DoGs are also generated under other types of stress. To explore this, we compared the DoGs recently reported to be generated in response to hyperosmotic stress in HEK293T cells [Rosa-Mercado et al., 2021] and those observed in breast cancer and pancreatic cells after TPL and THZ1 treatment. Out of the 606 DoGs generated in breast cancer cells in response to transcriptional stress, only 41 overlapped with those reported in cells subjected to osmotic stress (Fig. 4A). Similarly, in pancreatic tumor cells treated with TPL and THZ1, only a subset of the DoGs was shared with those induced by hyperosmotic stress (Fig.4C and Fig. 4E). This indicates that different types of stress can induce the formation of DoGs from different genes, although this phenomenon may also depend on the cell type (see discussion).

**Fig. 4.**
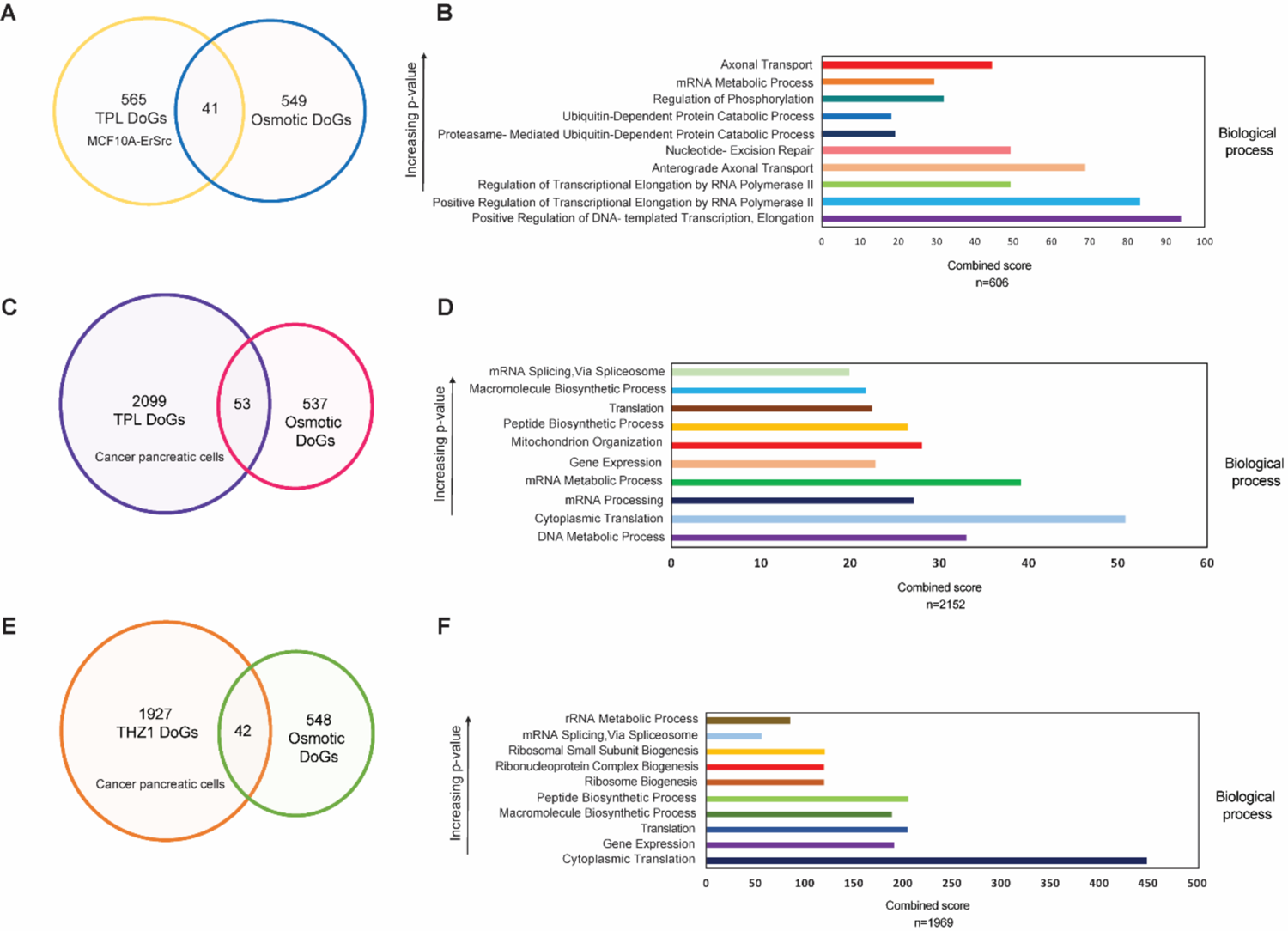
DoGs generated in response to transcriptional stress are mostly different from those generated by osmotic stress and are not related to a specific cell function. A) Venn diagram showing that of the 606 DoGs generated by transcriptional stress in cancer breast cells, only 41 are also produced by osmotic stress. B) Ontological analysis showing the biological functions of genes in which DoGs are generated by transcriptional stress in the breast cancer cells. C) Venn diagram showing that only 53 DoGs are shared between the cancer pancreas cells generated by TPL treatment and the produced by produced by osmotic stress. D) Ontological analysis showing the biological functions in which genes in which DoGs are generated by TPL treatment in pancreatic cancer cells. E) Venn diagram showing that only 42 DoGs are shared between the cancer pancreas cells generated by THZ1 treatment and the produced by produced by osmotic stress. F) Ontological analysis showing the biological functions in which genes in which DoGs are generated by THZ1 treatment in pancreatic cancer cells.

We also sought to determine whether particular types of genes with specific functions are more likely to generate DoGs. To this end, we performed ontological analysis of the DoGs generated in the breast cancer cell line and the pancreatic cancer cells (Fig. 4B, Fig. 4D and Fig. 4F). While the ontological analysis revealed a wide effect on different molecular functions of the genes that produce DoGs in the TSR, it is intriguing that many of these genes are related to processes that involve the synthesis of RNA (Fig 4B, Fig. 4D and Fig. 4F). However, at this point, it is not possible to correlate the genes that generate DoGs in the TSR with a specific function. In addition, we performed a GO analysis on the 668 DoGs shared by pancreatic cells treated with TPL and THZ1, as well as on the 67 DoGs found in the 3 conditions analyzed (Sup Fig. 2). Although the 668 genes shared by cancer pancreatic cells treated with the two drugs tend to be related to molecular functions associated with gene expression, the 67 DoGs found in the 3 conditions do not show a clear correlation with a specific cellular function.

## Discussion

The response to different types of stress is fundamental for the survival of cells when they are subjected to different types of insults. On the other hand, the stress response can also determine whether it is preferable for a cell to die when the damage is very severe. In general, any insult that generates stress in a cell causes the inactivation and overexpression of genes. A characteristic observed under oxidative stress, heat shock and viral infections is the generation of DoGs [Shalgi, 2014; Cugusi. et al., 2022; Heinz et al., 2018; Morgan 2022; Nemeroff 1998; Rosa-Mercado 2021; Rutkowski, et al., 2015]. We recently described that stress due to inhibition of the initiation of transcription mediated by RNA pol II causes the overexpression of some genes [Uriostegui-Arcos et al., 2020]. With the analysis presented here, we show that the TSR in breast cancer cells also leads to the generation of DoGs. This reinforces that after inhibition of transcription initiation using substances that inhibit the enzymatic activities of the TFIIH complex components, the cellular response is similar to that after other types of insults.

Intriguingly, we did not find a correlation between genes that are overexpressed in response to transcription inhibition and those that generate DoGs. This suggests that the mechanism by which some transcripts are extended in response to stress does not depend on an increase in transcription. We also found that the DoGs generated in response to TPL treatment are not exclusive to breast cancer cells but are also generated in cells derived from pancreatic cancer. Interestingly, a significant number of genes from which DoGs are generated during the TSR are shared between breast and pancreatic cancer cells treated with TPL. This suggests that there are genes that, either due to their function or due to the characteristics of their location in the genome, are more prone to DoG generation. However, by ontological analysis, we found that there does not seem to be a correlation among the functions of these genes, and when these genes are compared with genes from which DoGs are generated in response to hyperosmotic stress, there is not a strong correlation. This is a pattern that other groups have observed during investigations of different types of stress in different cell types [Morgan,et al., 2022; Rosa-Mercado et al., 2021; Rosa-Mercado and Steitz 2022]. In addition, we found that an important number of DoGs generated by TPL and THZ1 in pancreatic cancer cells are unique for each treatment, even that both substances are inhibitors of transcription. These could be related to the fact that TPL is a more severe drug than THZ1 and therefore the generation of some DoGs different. Also, side effects cańt be discarded, however more research is needed to answer these discrepancies.

A common feature when transcription is affected, is the degradation of the RNA pol II large subunit, as it is the case when TFIIH is affected (Chen et al., 2015; Uriostegui-Arcos et al., 2020). Thus, other substances that affect the PIC components may also generate DoGs. Therefore, it will be relevant in future analysis to determine if the different inhibitors of the RNA pol II also generate DoGs. It is possible that the inhibition of transcription has an impact effect in the factors that participate in transcription. Recent studies in cells exposed to osmotic stress have shown that the Integrator catalytic subunit Int11 which is required for the correct termination of many transcripts, the hyperosmotic stress induce its ubiquitination and in consequence its depletion, resulting in DoG generation [Rosa-Mercado et al., 2021]. It would be interesting in future studies to determine if the Integrator complex is also affected during the TSR and if it is also the cause of DoGs formation. On other hand, the question if the mechanism in the generation of DoGs is the same in different cellular stress conditions remains open, as well if in all cases the integrator is involved or if other factors are also involved.

The transcripts that form DoGs remain sequestered in the nucleus and are therefore not translated [Morgan et al., 2022; Rosa-Mercado and Steitz 2022]. Therefore, it has been proposed that their retention in the nucleus may be part of a stress protection mechanism to maintain chromatin integrity [Vilborg et al., 2015]. If so, the observation that RNA pol II extends the transcription elongation of a gene to regions that are not normally transcribed implies that the chromatin in those sites must be more open. Whether this renders the chromatin more protected against or more sensitive to damage is undetermined. On the other hand, the fact that mRNA with a DoG remains in the nucleus and therefore it cańt be translated, could be an alternative mechanism to inhibit the production of proteins when the cell in under stress. The fact that DoGs are generated during viral infections supports this hypothesis [Hennig et al., 2015].

In conclusion, inhibition of transcription generates a typical stress response in cells, in which not only is the overexpression of certain genes is induced but, as under other types of cellular stress, DoGs are generated. These two types of responses are particularly important for the treatment of cancer cells with transcription inhibitors since their effects may be involved in the generation of cells resistant to treatment.

## Materials and Methods

RNA-seq raw data was downloaded from NCBI using SRA-tools, for breast cancer cells GEO accession: GSE135256 and RNA-seq_T-1, RNA-seq_T-2, RNA-seq_TT-1, RNA-seq_TT-2 samples were used (GEO accession: GSM3999751, GSM3999752, GSM3999753, GSM3999754); for pancreatic cancer cells GEO accession: GSE157927 and Pancreatic cancer cells - Vehicle (DMSO) treated RNA-Seq Replicate 1, Pancreatic cancer cells - Vehicle (DMSO) treated RNA-Seq Replicate 2, Pancreatic cancer cells - Triptolide treated RNA-Seq Replicate 1, Pancreatic cancer cells - Triptolide treated RNA-Seq Replicate 2, Pancreatic cancer cells - THZ1 treated RNA-Seq Replicate 1, Pancreatic cancer cells - THZ1 treated RNA-Seq Replicate 2 samples were used (GEO accession: GSM4781045, GSM4781046, GSM4781047, GSM4781048, GSM4781049, GSM4781050 respectively).

FastQC-v0.11.7 was used to assess the quality of the sequence data. RNA-seq reads were aligned to the hg19 reference genome using Bowtie2 v. 2.3.4.3 with default parameters.

Samtools v. 1.9 was utilized to generate BAM files for each experiment and replicate. To identify DoGs, BAM files from breast and pancreatic cancer cells were processed using ARTDeco ‘get_dogs’ mode. A GTF file from UCSC (hg19 from Ensembl) was used with default settings, including a minimum DoG length of 4kb, a DoG window size of 500 bp, and a minimum FPKM of 0.2 for DoG discovery.

### Filtering DoGs with ChIP-seq Data

DoGs BED files generated by ARTDeco for MCF10A-Er-Src cells were used. RNAP II ChIP-seq BED files containing promoter peak coordinates for both DMSO and TPL conditions were obtained from GEO accession GSE135256 (supplementary files GSE135256_TSS_InPeaks_Tc_unique.bed/GSE135256_TSS_InPeaks_TTc_unique.bed). Two filters were applied:

– Filter 1: Checked for peaks that mapped inside DoGs. If a DoG covered a downstream gene with an RNAP II peak, it was considered as potential transcription of the downstream gene and discarded.
– Filter 2: Looked for DoG-producing genes with an RNAP II peak in their promoters to validate their transcription status.

Both filters were implemented using Python scripts available on the GitHub project page as Filter_1_with_ChIPseq.py and Filter_2_with_ChIPseq.py.

### DoGs Count and Comparison

The number of DoGs for each condition, replicate, and cell line was determined by counting the lines in each BED file generated by ARTDeco. To identify common DoGs between replicates, conditions, or cell lines, a Python script named Common_DoGs.py was used. This script searched for similar Ensembl IDs assigned to DoGs in two different BED files. To merge two or more DoGs files, the Union_DoGs_annotation command from DoGFinder was employed [Wiesel et al., 2018]. Bar graphs and Venn diagrams were created using R for visualization.

### Statistical Analysis

ARTDeco ‘diff_exp_read_in’ and ‘diff_exp_dogs’ modes were used to obtain the log_2_ fold change and p-values from DoG-producing genes and DoGs, respectively. With the output of those files and the DoGs files mentioned above, the heatmaps and scatter plots were generated with Python scripts (the Pearson’s correlation coefficient showed in the scatter plots was calculated using the SciPy library) available on the GitHub project page as Heatmap_log2fc.py and Scatterplot.py. Boxplots were created using the modified DoGs files, including DoG length information, and executed with the Boxplot.py code from the GitHub project page.

### Comparison of Genes

Names of DoG-producing genes from osmotic stress were acquired from table S1 [Rosa-Mercado et al., 2021]. Ensembl IDs for DoGs from TPL and THZ1 treatments in both cell lines were extracted and converted using DAVID Bioinformatics Resources [Huand et al., 2009; Sherman et al., 2022] to official gene symbol. The Stress_comparison.py code was utilized to identify common genes between DoG-producing genes from TPL or THZ1 and those from osmotic stress.

### Gene Ontology Analysis

Gene ontology analysis of biological processes was conducted using EnrichR [Chen et al., 2013; Kuleshov, 2016].

### GitHub Project page for code and data

https://github.com/PacoRM24/Transcriptional_stress_response_and_readthrough_transcription

## Acknowledgments

We thank the help of Dr. Diego Cortez for suggestions in the bioinformatic analysis and the bioinformatic unit at the Institute of Biotechnology for the use of the cluster. This work was supported by a grant from the PAPIIT/UNAM no. IN200621 and PASPA fellowship to Mario Zurta.

## Abbreviations

DoG: downstream-of-gene

lncRNA: long non-coding RNA

TFIIH: transcription factor II H

TPL: triptolide

TSR: transcriptional stress response

PIC: pre-initiation complex.

## Notes

### Competing Interest Statement

The authors have declared no competing interest.

