## Supplementary material for "Transcriptional stress induces the generation of DoGs in cancer cells": Sup. Figures

A B C

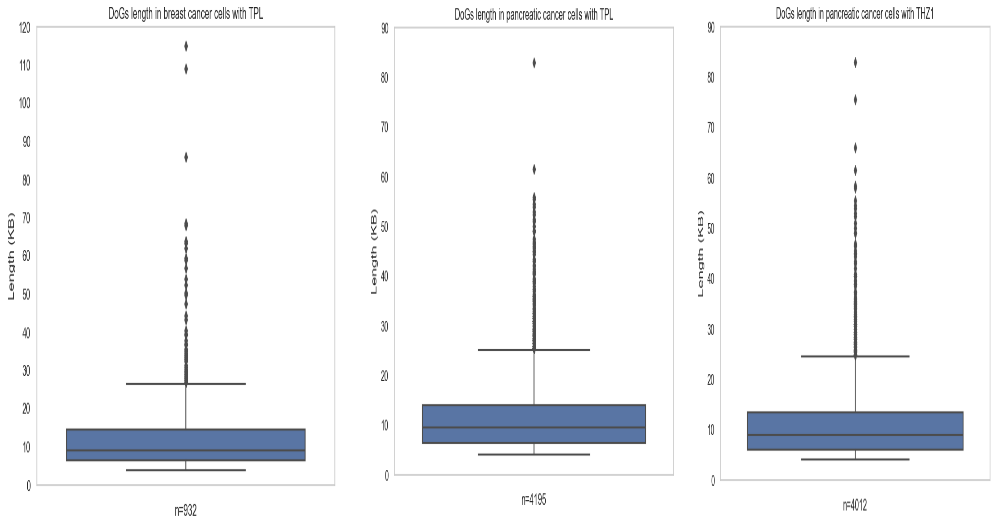


**Supplementary figure 1**. Box-plot representing the number of DoGs in relation to its length. A) DoGs generated in the breast cancer cell line in response to TPL treatment. B) DoGs generated in the pancreatic cancer cell line in response to TPL treatment. C) DoGs generated in the pancreatic cancer cell line in response to THZ1 treatment. Note that the median in the DoGs length is about 10,000 nucleotides in all cases.

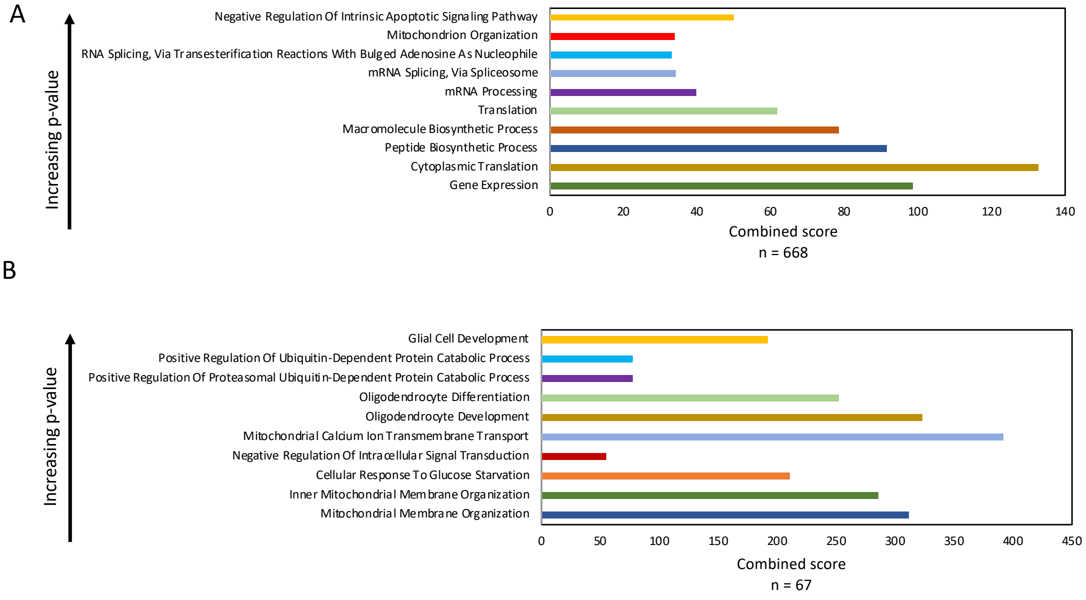


**Supplementary figure 2.** Gene Ontology Analysis. A) Gene ontology analysis of biological process from the 668 DoG-producing genes shared between pancreatic cancer cells treated with TPL and THZ1. B) Gene ontology analysis of biological process from the 67 DoG-producing genes shared between breast and pancreatic cancer cells treated with TPL and THZ1.
